# Binding Kinetics Oppositely Regulates type II Topoisomerase Relaxation and Decatenation Activities

**DOI:** 10.64898/2025.12.09.693212

**Authors:** Cleis Battaglia, Filippo Conforto, Yair Augusto Gutiérrez Fosado, Matt Newton, Erin Cutts, Davide Michieletto, Antonio Valdés

## Abstract

Type II Topoisomerases (topo II) are critical to simplify genome topology during transcription and replication. They identify topological problems and resolve them by passing a double-stranded DNA segment through a transient break in another segment. The precise mechanisms underpinning topo IIs ability to maintain a topologically simple genome are not fully understood. Here, we investigate how binding kinetics affects the resolution of two distinct forms of topological entanglement: decatenation and torsional relaxation. First, by single-molecule measurements, we quantify how monovalent cation concentration affects the dissociation rate of topo II from DNA. Second, we discover that increasing dissociation rates accelerate decatenation while slowing down relaxation catalytic activities. Finally, by using molecular dynamics simulations, we uncover that this opposite behaviour is due to a trade-off between search of target through facilitated diffusion and processivity of the enzyme in catenated versus super-coiled DNA. Thus, our findings reveal that a modulation of topo II binding kinetics can oppositely regulate its topological simplification activity, and in turn can have a significant impact *in vivo*.

---

Type II topoisomerases are a family of proteins involved in the topological regulation of genomes across life forms [1]. Type IIA topoisomerase (topo II) catalyses double-strand DNA passage through ATP-dependent process that involves temporary covalent bonds between its tyrosine residues and the 5’phosphate group in the backbone of a G-DNA segment, allowing a T-DNA segment to pass through [2–4]. The crucial role played by topo II is both well established and fascinating, as its ability to manage DNA entanglements appears essential for complex life [1]. However, the precise biophysical principles governing topo II–mediated topological simplification, especially in dense and crowded conditions of the nucleus or nucleoid, are poorly understood [5, 6]. While topo II can reduce the topological complexity of naked DNA below equilibrium in dilute conditions *in vitro* [7–9], it can also create catenated or knotted structures in dense conditions [10, 11] or in the presence of polycations [12]. Given the dense and entangled state of chromatin within the nucleus, how topo II maintains genomic simplicity *in vivo* remains unclear. Experiments on yeast minichromosomes [13] and human chromatin [14, 15] suggest that genomic topological complexity *in vivo* is lower than expected at equilibrium. Among the potential mechanisms to rationalise these observations are that topo IIs work in synergy with partner proteins, such as SMCs [6, 16, 17], or that they are post-translationally modified in order to change their binding and phase separation properties [18, 19]. The latter mechanism suggests a potential mode of regulation of topo II catalytic activity based on its binding kinetics [19, 20].

Binding kinetics is important for target search on coiled substrates like DNA [21], where proteins can perform diffusion in 1D, 3D and can perform intersegmental jumps (Fig. 1a). Faster binding and unbinding kinetics of topo II has been computationally suggested to yield faster topological simplification rates on knotted DNA substrates due to an effective “enhanced 3D sampling” of essential crossings [20]. However, it remains to be experimentally determined whether all catalytic activities of topo IIs are favoured by faster (un)binding kinetics. Motivated by this open question, in this work we tested topo II topological relaxation and decatenation activities on supercoiled and catenated DNA substrates to determine how they are affected by its (un)binding kinetics.

**FIG. 1.**
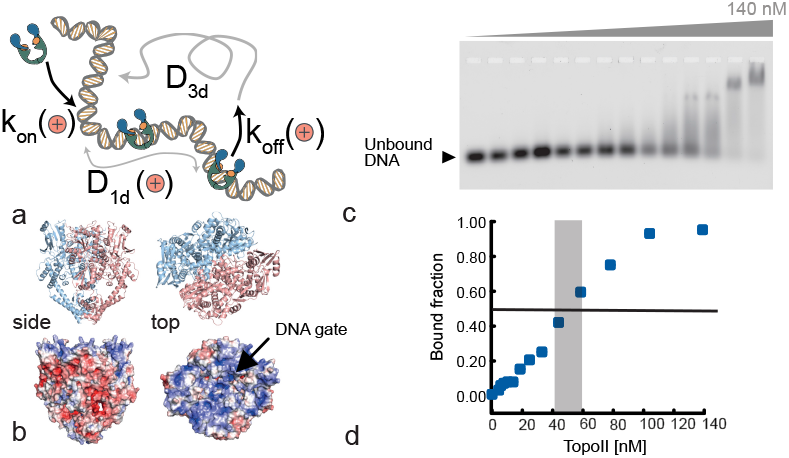
**a**. Topo II search on DNA, including 1D and 3D diffusion and intersegmental jumps. On and off rates, and 1D diffusion constant are expected to be salt dependent. **b**. Topo II present exposed positive surface charges that stabilise the binding with negatively charged DNA substrate. Its intrinsically disordered CTD also display positive charges that can bind to dsDNA (see SI). red = negative, blue = positive charges. **c**. Electrophoretic mobility shift assay (EMSA) performed on a 51 bp dsDNA varying topo II concentration from 0 to 140nM, at 125 mM NaCl. **d**. Quantification of the bound dsDNA fraction in the EMSA, returning an approximate *k*_*D*_ ≃ 50 nM.

First, we determined that topo II binding kinetics can be modulated by varying monovalent cations within a physiological range. By screening negative charges along the DNA, monovalent cations reduced the binding affinity of topo II to the DNA and in turn accelerated its unbinding rate. We thus leveraged this modulation and performed topological simplification assays on catenated and supercoiled structures in the presence of different amounts of monovalent cations. Surprisingly, we observed opposite effects: decatenation was accelerated by higher cation concentration, while relaxation was slowed down. We rationalised these observations by performing Molecular Dynamics simulations of topologically constrained or catenated DNA plasmids under the action of a dynamically binding topo II, mimicking alternating 1D and 3D diffusion. Our simulated results – in line with the experimental findings – suggest that whilst efficient decatenation benefits from fast binding kinetics, torsional relaxation is favoured by more processive topo II with longer binding times.

## RESULTS

### Topo II binding kinetics is modulated by monovalent cations

Monovalent cations change the binding affinity of DNA-binding proteins by screening electrostatic interactions. More specifically, they can significantly increase the dissociation rate (*k*_off_) of proteins from DNA whilst remaining within physiologically relevant concentrations (e.g., 50-150 mM NaCl) [22, 23]. Topo IIs display three principal positively charged surfaces that are thought to interact with, and stabilise, the nucleo-protein complex. A major patch lies along the DNA cleavage-rejoining domain, or DNA gate, which binds the G-DNA segment (Fig. 1b); and two additional patches located on the intrinsically disordered C-terminal domains (CTD); (Fig. S1a). We thus decided to quantify the (un)binding kinetics of yeast topo II on DNA for different physiological levels of monovalent cations (Na^+^). First, we precisely measured the dissociation kinetics *k*_off_ at single molecule resolution. We used confocal microscopy to visualise single fluorescently labelled topo II binding, diffusing and unbinding on *λ*DNA stretched using optical tweezers (see SI for details, Fig. 2a). We performed the same experiments at different levels of [Na^+^] and we then analysed the kymographs (over 100 tracks for each condition), extracted the length of the single traces, and finally determined the dwell time, or inverse dissociation constant *k*_off_ (Fig. 2b-e). The values of *k*_off_ measured range within 0.1 − 10 *s*^−1^ and significantly increase with higher concentrations of monovalent cations, in line with what observed for other DNA-binding proteins [24, 25]. We also measured the apparent dissociation constant *k*_*D*_ via electrophoretic mobility shift assay (EMSA) on a short (51 bp) dsDNA (Fig. 1c,d). We can compare these measurements with estimates from a Smoluchowski collision equation; the on-rate can be estimated as *k*_*on*_ ≃ 4*π*(*D*_1_ + *D*_2_)(*a*_1_ + *a*_2_) ≃ 5 × 10^9^*M*^−1^*s*^−1^ with *D*_1_ = *k*_*B*_*T/*(3*πη*_*w*_*a*_1_) ≃ 10*µm*^2^*/s, D*_2_ ≪ *D*_1_, *a*_1_ = 40 nm the size of topo II [26, 27], *a*_2_ = 10 nm the size of the target DNA site and where we used water viscosity *η*_*w*_ = 1 mPa s. This yields a dissociation constant *k*_o*ff*_ = *k*_*D*_*k*_*on*_ = 50*s*^−1^ which is within the same order of magnitude of the ones measured in single molecule experiments (Fig. 2f).

**FIG. 2.**
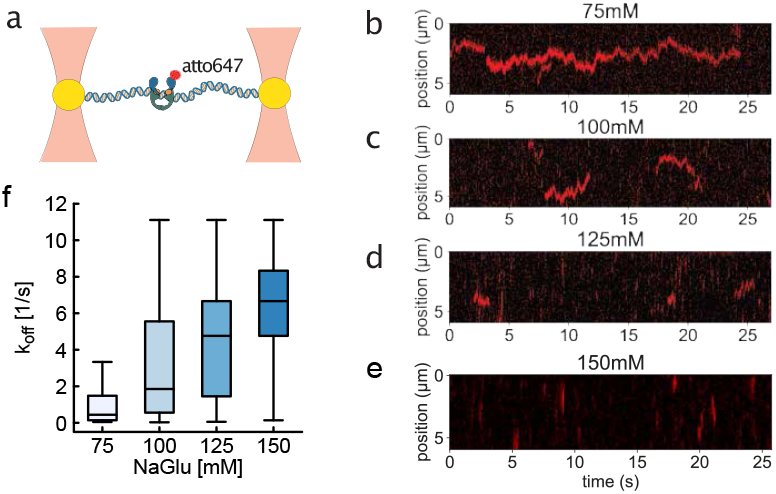
Binding kinetics of topo II is modulated by monovalent cations. **a**, Diagram of force-stretched DNA. DNA tethered between trapped beads, topo II complex is labeled using atto647. **c-f** Kymographs of atto647-labelled topo II binding, diffusing and unbinding from a stretched *λ*DNA using optical tweezers at 75-150 mM [NaGlu]. **g**. Quantification of inverse dwell time, or *k*_off_, as a function of monovalent cation concentration. Each concentration includes analysis of more than 100 single-molecule tracks. All pair-wise distributions are significantly different (p value < 0.01).

### Faster binding kinetics accelerates decatenation and slows down relaxation

Having quantified how the dwell time *τ*_*off*_ = 1*/k*_*off*_ changes with the concentration of monovalent cations in solution, we then asked how this parameter affected the catalytic activity of topo II to perform topological decatenation of catenated, kinetoplast DNA (kDNA) [28, 29] (Fig. 3a). To this end, we incubated *Crithidia fasciculata* kDNA, a structure made of thousands of catenated DNA rings [28, 29], with yeast topo II at various concentrations of monovalent cations. The enzyme concentration was set to be comparable to the substrate molarity, and reaction conditions were selected to ensure that decatenation did not reach full topological simplification; reactions were then stopped and the samples were run on a gel to separate catenated from single-circle topologies. We observe that the decatenation rate is sped up by higher levels of NaCl, corresponding to shorter dwell times of topo II on DNA (Fig. 3b-c). This result is consistent with previous numerical results, where different modes of topo II (un)binding to DNA substrates were found to enhance topo II unknotting activity [20].

**FIG. 3.**
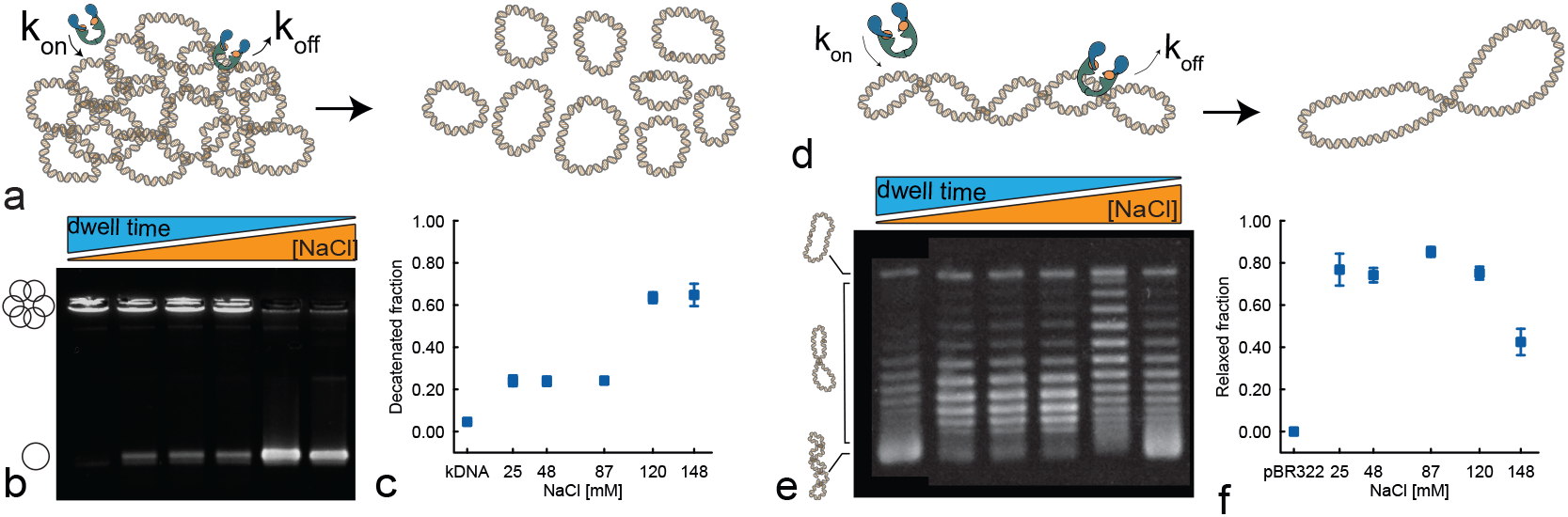
Topoisomerase decatenation and relaxation activities are oppositely regulated by salt concentration. **a**. To quantify decatenation activity we monitor the disassembly of kDNA as a function of various [NaCl]. **b**. Gel electrophoresis of 20 ng/ul kDNA after 30 minutes incubation with 3.45 ng/ul of topo II. The first lane shows the control sample without protein, and the following five lanes correspond to reactions performed in buffers containing [NaCl] = 25, 48, 87, 120, and 148 mM, respectively. **c**. Quantification of decatenated minicircles as a function of [NaCl]. **d**. To quantify the relaxation activity we monitor the topology of a negatively supercoiled pBR322 plasmid as a function of various [NaCl]. **e**. Chloroquine gel electrophoresis of 20 ng/ul pBR322 after 30 minutes incubation with 3.45 ng/ul of topo II. As in panel b, the first lane corresponds to the control sample without protein, and the remaining lanes show reactions carried out at [NaCl] = 25, 48, 87, 120, and 148 mM. Notably the amounts of supercoiled and relaxed DNA at 120mM and 87mM NaCl are comparable, nevertheless the supercoil distributions are visibly different, showing a slowed relaxation effect at higher NaCl concentrations. **f**. Quantification of relaxed topoisomer fraction as a function of [NaCl].

We then performed topological relaxation experiments where a 4.5 kb negatively supercoiled plasmid (pRB322) was incubated with yeast topo II and varying [NaCl] (Fig. 3d). As for the decatenation assay, the enzyme concentration was kept comparable to the substrate molarity, and conditions were chosen to prevent the reaction from reaching full topological relaxation. We then ran the samples through a gel in presence of 0.6 *µ*g/ml of chloroquine and quantified the distribution of topoisomers (Fig. 3e-f). The enzyme concentration was set to be comparable to the substrate molarity, while reaction conditions were selected to ensure that, given the enzyme’s activity, neither decatenation nor relaxation proceeded to full topological simplification. This allowed us to resolve intermediate topological states and evaluate the salt de-pendence of activity. Unexpectedly, we found that topo II was less efficient at relaxing supercoiling at larger salt concentrations. In fact, while at low [NaCl] the topoisomer distributions were tightly peaked around the relaxed *Lk*_0_, at larger [NaCl] we could observe a very broad distribution (5th lane in Fig. 3b). This observation is consistent with the short dwell time of topo II on DNA and suggests that the enzyme performs short rounds of catalytic activity in these conditions. On the other hand, it is conceptually opposite to what we observed in the decatenation experiment. These results were consistent in both sodium chloride and sodium glutamate and did not depend on the presence of the CTD (Fig. S6), even though the CTD is a strong DNA-binding site (Fig. S1b).

### MD simulations explain the opposite catalytic regulation by monovalent cations

To rationalise our experiments, we decided to perform Molecular Dynamics (MD) simulations of a coarse-grained bead-spring polymer model of DNA under the action of kinetically binding topo II.

First, we simulated a 1 *µ*m^2^ patch of kinetoplast DNA made of 90 linked DNA minicircles, each made of 2.5 kb or 85 beads, whose topology was inferred from AFM data (Ref. [29]). We then run a Langevin simulation (implicit solvent, see SI) including a sub-stoichiometric number of topo IIs (*N*_*t*_ = 80), which were modelled by turning a 5-beads segment of the polymer into a “phantom” seg-ment. During the simulation, the polymer chains undergo Brownian motion and most of the beads interact via a uncrossable, purely repuslive potential. The *N*_*t*_ = 80 “phantom” segments representing topo II are the only segments that can be freely crossed and are transiently added/removed from the polymers stochastically, with typical on- and off-rates that mimic different [NaCl] conditions (Fig. 4a). More specifically, we fixed the on-rate to 0.001 1*/τ*_*B*_ and increased the off-rate from 10^−4^1*/τ*_*B*_ to 1*/τ*_*B*_ to capture shorter dwell times at increasing salt concentrations. Additionally, we assume that topo II does not perform 1D diffusion along the rings. During the simulation we track the topology of the network via the pairwise linking number

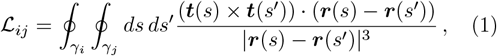

where *γ*_1_ and *γ*_2_ are simulated polymer rings, ***t***(*s*) and ***r***(*s*) are the tangent and position of the polymers at arclength *s*. Given ℒ_*ij*_ for all the pairs of rings, we can calculate how many rings belong to the largest interlocked component in the system. Because we start from a fully catenated structure, the fraction of catenated rings *f*_*C*_ will start from one and decay in time. By averaging over 50 independent replicas of the system we thus obtain the average fraction of catenated ⟨*f*_*C*_(*t*) ⟩ plotted in Fig. 4b. These curves can be used to directly compare our simulations with the gels in Fig. 3b-c. Specifically, we compare ⟨*f*_*C*_⟩ at fixed runtime 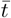 and for different values of *k*_off_ and observe that the larger the off-rate the larger the smaller the catenated fraction – and hence the larger the decatenated fraction – as seen in experiments. Interestingly, and in line with experiments, we also observe that 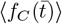 plateaus at large *k*_off_ suggesting that the decatenation process is reaction limited, i.e. a minimum topo II dwell time is required to perform successful decatenation. In other words, it is not possible to accelerate the decatenation process indefinitely after some value of the off-rate, which is required to catalyse at least one double strand passage.

**FIG. 4.**
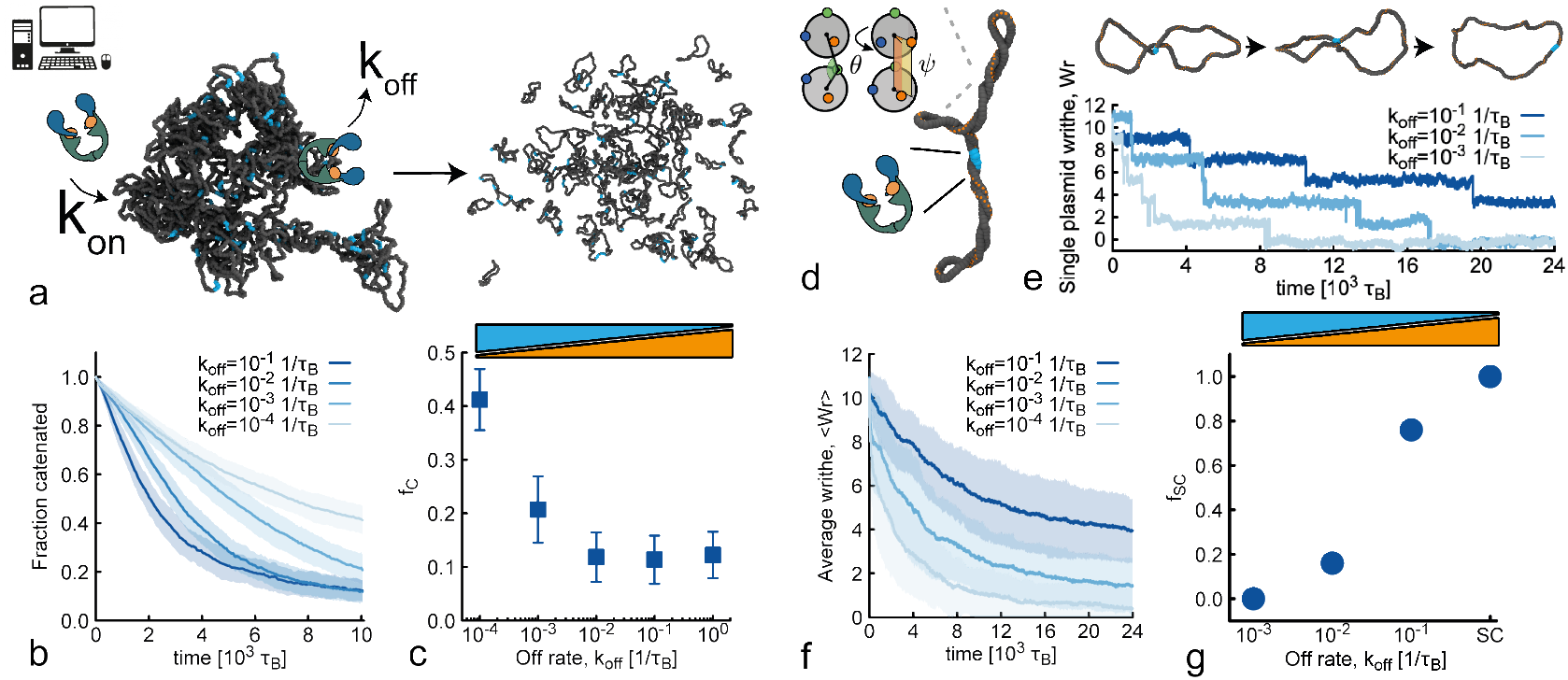
Molecular Dynamics simulations with salt-dependent *k*_off_ capture the opposite topological regulation. **a**. Snapshots from MD simulations of a model kDNA from Ref. [29] before and after decatenation reaction. **b**. Fraction of catenated minicircles as a function of time and for different values of off-rate. Shaded area represents the standard deviation over 50 independent replicas. **c**. Fraction of catenated minicircles at fixed time (10^4^*τ*_*B*_) and as a function of *k*_off_ showing enhanced decatenation at larger off-rate. **d**. Snapshot from MD simulations of a supercoiled DNA chain (model from Ref. [30, 31]. **e**. Step-wise change in writhe monitored through the simulation. Snapshots at the top, plot of writhe at the bottom. **f**. Average writhe ⟨𝒲⟩ as a function of time and for different values of off-rate. The shaded area represents the standard deviation across 50 independent replicas. **g**. Fraction of supercoiled topoisomers (defined as those with 𝒲*/N >* 0.05) as a function of off-rate (SC = fully supercoiled state).

Having observed that our simulations can recapitulate the experimental results for the decatenation activity of topo II we decided to computationally test its relaxation activity. To model supercoiled DNA we employ a coarse-grained “twistable” polymer (see Refs. [30, 32] for details). We implement a dynamically binding/unbinding topo II by transiently creating/deleting a “phantom” segment from the chain (see Fig. 4a and Methods). We maintain the on-rate constant and increase *k*_off_ to capture the faster unbinding at larger [Na^+^]. The topological relaxation of our supercoiled polymer can be quantified by measuring its writhe [33, 34]

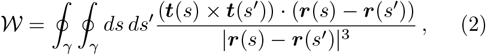

where ***t***(*s*) and ***r***(*s*) are the tangent and position of the curve *γ* at arclength *s*. The relaxation proceeds by allowing the polymer backbone to cross through itself, as shown in Fig. 4b. Each strand-crossing event yields a step-wise change in writhe Δ𝒲 = −2. Note that the writhe is not an integer and can fluctuate as long as the sum of twist and writhe of single chains remain constant (CFW theorem [35]). By averaging over 50 independent replicas we then obtain relaxation curves ⟨𝒲 (*t*)⟩ plotted in Fig. 4f. Surprisingly, and opposite to what we saw for the decatenation reaction, we observe that larger *k*_off_ leads to slower relaxation (Fig. 4g). However, this slowing down is consistent with the experiments in Fig. 3 and demonstrates opposite regulation of catalytic activities of topo II simply by a change in monovalent cation concentration.

## DISCUSSION

In this work we observed a significant and opposite regulation of topo II topological decatenation and relaxation activities as a function of monovalent cation concentration.

We rationalise the observed behaviour as follows: while at large [Na^+^] topo II spends more time performing unbound 3D search, at small [Na^+^] topo II spends more time bound to the DNA. Thus, in the regime of low monovalent cation concentration, topo II is highly processive and performs several rounds of strand-passage without leaving the substrate. In contrast, at high monovalent cation concentration topo II is less processive, quickly dissociating from DNA and spending more time performing facilitated 3D diffusion.

These different binding kinetics have profound effects on topo II catalytic activities. Supercoiled plasmids have highly localised entanglements and are therefore rapidly relaxed by a highly processive enzyme that remains bound long enough to perform sequential strand-passage events. On the contrary, entanglements resulting from catenation are typically delocalised across the substrate and therefore too long binding times yield inefficient decatenation because after a strand passage, no further entanglements are present at a given location. Thus, efficient decatenation is achieved through rapid binding–unbinding dynamics that allow topo II to disengage quickly and relocate to remaining entanglements. Our results thus reveal an intrinsic kinetic trade-off: torsional relaxation is favoured by prolonged binding, whereas decatenation is accelerated by faster turnover.

We argue that this opposite regulation may have physiological relevance. Indeed, tuning the binding affinity of topo II to DNA, for example by post-translational modifications [19], it is possible to oppositely regulate topo II decatenation and relaxation activities.

This finding is consistent with the view that topoisomerases are recruited and controlled through pro-tein–protein interactions and post-translational modifications that localize them to specific genomic regions or cell-cycle stages. For example, recent studies show that the oncoprotein MYC can form a complex with human topo II*α*, enhancing topoisomerase retention at regions experiencing elevated transcription-induced supercoiling [36, 37]. In this framework, increasing enzyme dwell time at highly supercoiled loci would promote efficient relaxation, consistent with our observations under low-salt conditions. Conversely, during stages of the cell cycle that demand rapid decatenation, such as metaphase, when sister chromatids must be fully separated before anaphase, excessively long enzyme dwell times could be detrimental, increasing the likelihood of unwanted re-catenation events. Our results under high-salt conditions, which shorten DNA residence time and accelerate decatenation, suggest a biophysical basis for how cells might avoid such outcomes. Together, our findings provide mechanistic insight into how topo II activity can be tuned through its binding kinetics, offering a unified explanation for how cells could differentially regulate supercoil relaxation and chromosome decatenation. By demonstrating how subtle shifts in dwell time produce qualitatively distinct topological outcomes, this work highlights binding kinetics as a central and under-appreciated layer of topo II regulation *in vivo*.

## MATERIALS AND METHODS

### Protein expression and purification

Full-length and C-terminal domain truncated topo II were expressed in yeast (see SI for detailed expression and purification protocol). Ybbr peptide tag on full-length topo II was labelled with Sfp phosphopantetheinyl transferase (P9302, New England Biolabs) and CoA-ATTO647N (see SI for detailed fluorescent labelling protocol).

### Electrophoretic mobility assay

Electrophoretic mobility assay (EMSA) was carried out by mixing a 6-FAM labelled 51 bp dsDNA with protein at the concetrations indicated in the figure in 40 mM TRIS–HCl pH 7.5, 125 mM NaCl, 5 mM MgCl2, 10% glycerol, 1 mM DTT. After 10 min incubation at room temperature ( 25° C), free DNA and DNA–protein complexes were resolved by electrophoresis for 1 h at 4 V/cm on 0.75% (w/v) TAE-agarose gels at 4°C. We then imaged the 6-FAM labelled DNA using a gel scanner (see SI for more details).

### DNA relaxation assay

200 ng of pBR322 (NEB) plasmid were incubated with 34.5 ng of topoII-ΔCTD in a 10 ul volume containing 50mM Tris–HCl (pH 7.5), [25,48,87,120,148] mM NaCl, 5 mM MgCl2, 1 mM DTT. ATP was added to 1mM and the mixture was incubated at 37°C during 30 min. Reactions were stopped with the addition of 20mM EDTA, 1% (w/v) sodium dodecyl sulphate (SDS). Subsequently, 0.8U of proteinase K were added, followed by incubation for 1 h at 65°C. Reaction samples were loaded onto 1% (w/v) agarose gels. DNA electrophoresis was performed at 2 V/cm for 15 h in 1X TAE buffer (40 mM Tris, 20 mM acetate, 1 mM EDTA) containing 0.6 ug/mL chloroquine. Gels were then stained with SYBR Gold (Thermo Fisher Scientific) and visualized using a Syngene transilluminator. Gel images were analyzed using Fiji [38] (see SI).

### DNA decatenation assay

The protocol used for the decatenation assay is the same as the one used for the relaxation assay, except for the following differences: Instead of pBR322 plasmid, 200 ng of Kinetoplast DNA (Inspiralis) are added to the reactions. No chloroquine is used for the electrophoresis.

### Molecular Dynamics simulations

Molecular Dynamics simulations were performed in LAMMPS [39] in NVT ensemble using a Langevin thermal bath and custom fixes that changed the type of beads during the course of the simulations. In both relaxation and decatenation simulations we considered sub-stoichiometric number of topo IIs, in line with experiments. Topo IIs were modelled by turning a 5-beads segment of the polymer into a “phantom” segment. All other beads along the polymers interacted via a uncrossable, purely repulsive potential. The “phantom” segments were transiently added/removed from the polymers stochastically, with typical on- and off-rates that mimic different [NaCl] conditions (Fig. 4a). More specifically, we fixed the on-rate to 0.001 1*/τ*_*B*_ and increased the off-rate from 10^−4^1*/τ*_*B*_ to 1*/τ*_*B*_ to capture shorter dwell times at increasing salt concentrations. See SI for more details of the simulations.

### Single molecule assay

Single molecule assay was conducted using the Lumicks C-Trap instrument BA105, equipped with integrated confocal microscopy and microfluidics (Fig. 2a, SI Fig.2a). The fluidics was cleaned by flowing 500 ul 2% Hellmanex over 40 minutes followed by flowing 2 x 0.5 ml MQ H2O and finally 0.5 ml Optical tweezers Buffer, OTB (50mM Tris–HCl (pH 7.5), [75,100,150,200] mM NaGlu, 5 mM MgCl2, 1 mM DTT). Protein channels (Channels 4 and 5) were passivated with 0.5 ml Casein (0.01% w/v) and BSA (0.005% w/v) in OTB flowed through over a 30 minute period. *λ*-DNA was prepared by ligation of biotinylated caps (as previously described [40, 41]. Fluidics channels were prepared as follows: Channel 1, 0.005% w/v 4.38 um SPHERO™Streptavidin coated polystyrene beads (SVP-40-5) in OTB; Channel 2, biotinylated DNA diluted 200-fold in OTB; Channel 3, OTB; Channels 4 and 5, topo2-atto647 at 2-3 nM in OTB. OTB in Channels 4 and 5 was prepared with varying NaGlu concentrations (75, 100, 150, or 200 mM) while kept at constant NaGlu concentration of 100 mM in the other channels. DNA was and tethered between optically trapped beads using the laminar flow cell. Prior to each experiment, a force-extension curve was measured from 0–55 pN to confirm the presence of a single intact DNA molecule. DNA was held at 5 pN before moving into the channel containing protein. Confocal imaging was performed using 5% excitation power at 647 nm, with emission detected through a 680/42 nm band-pass filter; images were acquired with a pixel size of 100 nm and a pixel dwell time of 0.1 ms, using a line width of 25.5 *µ*m and a line time of 30 ms (See SI for data analysis).

## DATA AVAILABILITY

All the information are included in the Supplementary Information. Codes are deposited at https://git.ecdf.ed.ac.uk/taplab.

## ACKNOWLEDGEMENTS

D.M. acknowledges the Royal Society and the European Research Council (grant agreement No 947918, TAP) for funding. We thank Nesibe Durmaz for technical assistance. The authors also acknowledge the contribution of the COST Action Eutopia, CA17139. For the purpose of open access, the author has applied a Creative Commons Attribution (CC BY) licence to any Author Accepted Manuscript version arising from this submission. M.D.N. is supported by a Wellcome Early Career Award (225139/Z/22/Z). The LUMICKS C-trap instrument was funded through MRC Equipment Grant MC PC APP28599 and The Henry Royce Institute for Advanced Materials, funded through EPSRC grants EP/R00661X/1, EP/S019367/1, EP/P02470X/1 and EP/P025285/1. The work was funded by Royce Access Scheme SHEF-YR10-003 SREAS25/006. We would like to acknowledge Xinyue Chen for technical support at Royce@Sheffield.

## MATERIAL AND METHODS

### Protein expression and purification

Ybbr tagged full-length, and C-terminal domain truncated (Δ1196 − 1428) of *Saccharomyces cerevisiae* topoisomerase II (topoII-ΔCTD) were expressed in yeast from 2*µ*-based high copy plasmids under the control of galactose-inducible promoter pGAL1. Cells were initially grown at 30°C –URA dropout media containing glucose to OD600 of 1, transferred to media containing 2% raffinose for 2 hours and then induced by addition of galactose to 2% for 8 hours. Cells were harvested by centrifugation, re-suspended in buffer A (50 mM TRIS-HCl pH 7.5, 300 mM NaCl, 5% (v/v) glycerol, 5 mM *β*-mercaptoethanol, and 20 mM imidazole) containing 1X complete EDTA-free protease-inhibitor mix (Roche) and lysed in a FreezerMill (Spex). The lysate was cleared by two rounds of 20 min centrifugation at 45000g at 4°C and loaded onto a 5 ml HisTrap column (GE Healthcare) preequilibrated with buffer A. The resin was washed with five cv buffer A; buffer A containing 1 mM ATP, 10 mM KCl and 1 mM MgCl2. Protein was eluted in buffer A containing 300 mM imidazole. The buffer of the eluate fractions was exchanged to low salt buffer (25 mM Tris-HCl pH 7.5, 150 mM NaCl, 5% (v/v) glycerol, 1 mM DTT) using a desalting column before loading onto a RESOURCE Q anion exchange column (Cytiva) preequilibrated with low salt buffer. After washing with 3 cv low salt buffer, proteins were eluted by increasing NaCl concentrations to 1 M in a linear gradient of 60 mL. Peak fractions were pooled and concentrated by ultrafiltration (Vivaspin 30,000 MWCO, Sartorius) before loading onto a Superose 6 increase 10/300 column (Cytiva) pre-equilibrated in buffer B (25 mM Tris-HCl pH 7.5, 300 mM NaCl, 5% (v/v) glycerol, 1 mM DTT). Peak elution fractions were concentrated by ultrafiltration, snap-frozen and stored at –80°C until use.

The yeast topoisomerase II CTD (G1178-D1428) (topo IICTD) construct with an N-terminal His8-mNeonGreen tag was expressed in *E. coli* Rosetta (DE3) pLysS (Merck) grown at 37 °C in 2× TY medium. Protein expression was induced with IPTG at a final concentration of 0.2 mM and cultures were shifted to 18 °C. Cells were collected 18 h post-induction by centrifugation, snap frozen in liquid nitrogen, and stored at –80 ºC. Cells were thawed and lysed by sonication at 4 °C in lysis buffer (50 mM TRIS-HCl pH 7.5, 200 mM NaCl, 20 mM imidazole, 1 mM *β*-mercaptoethanol) containing cOm–EDTA. The lysate was cleared by centrifugation at 48000 ×g for 30 min at 4 °C. The supernatant was incubated for 2–3 h with 2 mL Ni Sepharose HP (Cytiva) at 4 °C and eluted with elution buffer (50 mM TRIS-HCl pH 7.5, 200 mM NaCl, 250 mM imidazole). The eluate was dialyzed overnight against low-salt buffer (50 mM TRIS-HCl pH 7.5, 100 mM NaCl, 1 mM DTT) at 4 °C and loaded onto a 6-mL Resource S ion-exchange column (Cytiva) pre-equilibrated with low-salt buffer. After washing with 5 cv low-salt buffer, protein were eluted by increasing NaCl concentrations to 1 M in a linear gradient of 20 cv. Peak fractions were pooled and loaded onto a Superdex 200 increase GL 10/300 column (Cytiva) pre-equilibrated in SEC-buffer (50 mM TRIS-HCl pH 7.5, 300 mM NaCl, 1 mM DTT). Peak fractions were pooled and concentrated by ultrafiltration (Vivaspin 30000 MWCO, Sartorius) at 4 °C. The concentrated protein was aliquoted, snap frozen in liquid nitrogen, and stored at –80 °C.

### Fluorescent labeling of full-length yeast topoisomerase

Enzymatic site-specific covalent labeling of the Serine hydroxyl of ybbR peptide tags (GTDSLEFIASKLA) on full-length topo II (12 *µ*M) was performed in buffer B supplemented with 10 mM MgCl2, 1 *µ*M Sfp phosphopantetheinyl transferase (P9302, New England Biolabs) and 15 *µ*M CoA-ATTO647N conjugate reaction mix. After 16 h incubation at 4 ºC, topo II was purified by size-exclusion chromatography on a Superose 6 increase 3.2/300 column (Cytiva) preequilibrated in buffer B containing 1 mM MgCl2. Peak elution fractions were concentrated by ultrafiltration, snap-frozen and stored at –80 ºC until use.

### Electrophoretic mobility assay

6-carboxyfluorescein (6-FAM) labelled 51 bp dsDNA was prepared by annealing two complementary HPLC-purified DNA oligonucleotides (Merk, 5’-6-FAM-TAT TTT CTT GTT CTT TTA CAT CAA CAC AAT GTG ACC ATT ACC TGG TTA TCA-6-FAM-3’; 5’-6-FAM-TGA TAA CCA GGT AAT GGT CAC ATT GTG TTG ATG TAA AAG AAC AAG AAA ATA-3’) in annealing buffer (10 mM TRIS-HCl pH 7.5, 50 mM NaCl, 1 mM EDTA) at a concentration of 50 *µ*M in a temperature gradient of 0.1^°^C/s from 95^°^C to 4^°^C. 20 *µ*l reactions were prepared with 10 nM 6-FAM-dsDNA and protein concentrations ranging from 0 to X nM (as specified in captions) in reaction buffer (40 mM TRIS–HCl pH 7.5, 125 mM NaCl, 5 mM MgCl2, 10% glycerol, 1 mM DTT). After 10 min incubation at room temperature (∼ 25^°^C), free DNA and DNA–protein complexes were resolved by electrophoresis for 1 h at 4 V/cm on 0.75% (w/v) TAE-agarose gels at 4^°^C. 6-FAM labelled DNA was detected on a Typhoon RGB scanner (GE Healthcare) with excitation at 488 nm and detection using a Cy2 filter.

**FIG. 1.**
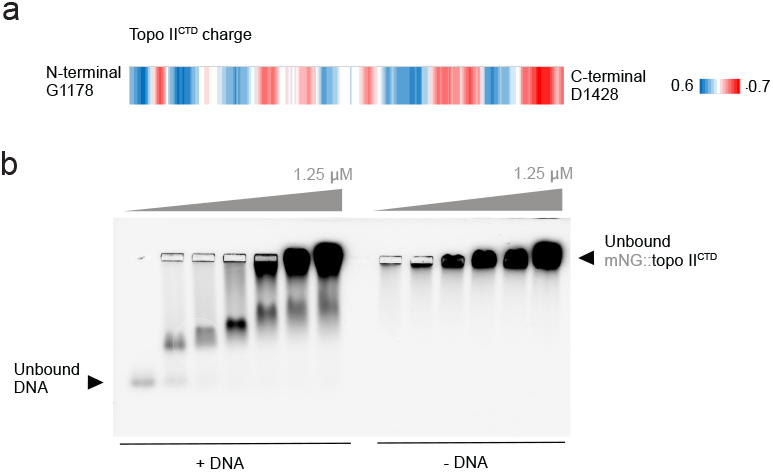
Yeast topo IICTD binds dsDNA. **a**. Charge distribution along the CTD of yeast topo II (blue positive, red negative, sliding window 10). **b**. Electrophoretic Mobility Shift Assay (EMSA) with 51 bp dsDNA and mNeonGreen tagged yeast topo II CTD (mNG::topo IICTD).

### Kinetic and binding distribution analysis

For kinetic analysis, binding events were identified from kymographs using the greedy algorithm implemented in Pylake software [1] using the following parameters: track width=0.6, pixel threshold=1.9, window=4, adjacency filter=True, min length=4, sigma=0.3, velocity=0, diffusion=1.5, sigma cutoff=1.5, use widgets=True, correct origin=True. Tracks shorter than four frames or intersecting each other or the beads were discarded. Tracks with intensities exceeding three standard deviations from the typical signal were excluded to restrict analysis to single-protein events. Detected binding events were analyzed using trackpy. To validate the tracking procedure, we applied it to multiple kymographs recorded in the absence of protein, where, as expected, no tracks were detected (see SI Fig.2b). Dwell times for each state were extracted from kymographs using a Gaussian Mixture Model (GMM) model. A total of 161, 137, 177, and 108 tracks were analyzed for the 75 mM, 100 mM, 125 mM, and 150 mM conditions, respectively. The distributions of observed dwell times were fitted using maximum likelihood estimation (MLE) with the DwelltimeModel class. Confidence intervals were estimated via bootstrapping as shown if SI Fig.3. Because dwell times at 150 mM NaGlu are close to the temporal resolution limit of the scans, this condition provides only an upper limit of the true dwell time. Notably, to minimize detection errors that occur when multiple binding events occur in close proximity, low 3 nM concentrations of topo2-atto647 for the [75,150,200]mM NaGlu and 2nM for 50mM NaGlu.

**FIG. 2.**
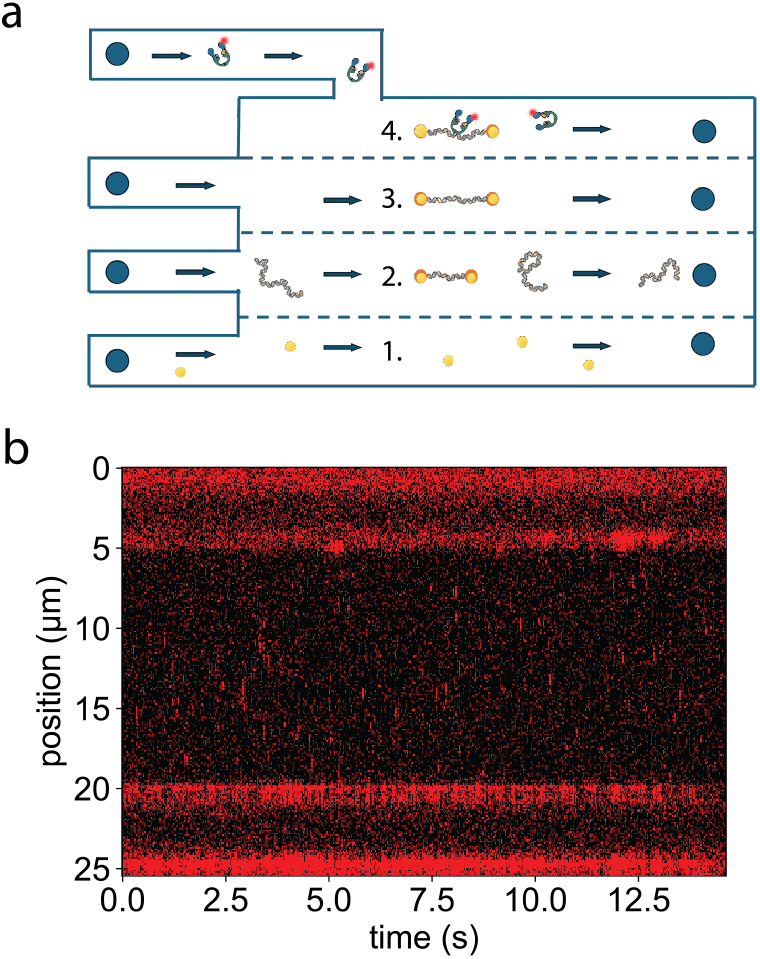
Microfluidic set-up and control kymograph in the absence of protein. **a**. Microfluidic set-up: 1. bead channel, 2. DNA channel, 3. buffer-only channel, 4. imaging channel. **b**. Representative kymograph acquired in the absence of protein, showing no detectable tracks. This confirms that the tracking algorithm does not produce false positives under these conditions.

**FIG. 3.**
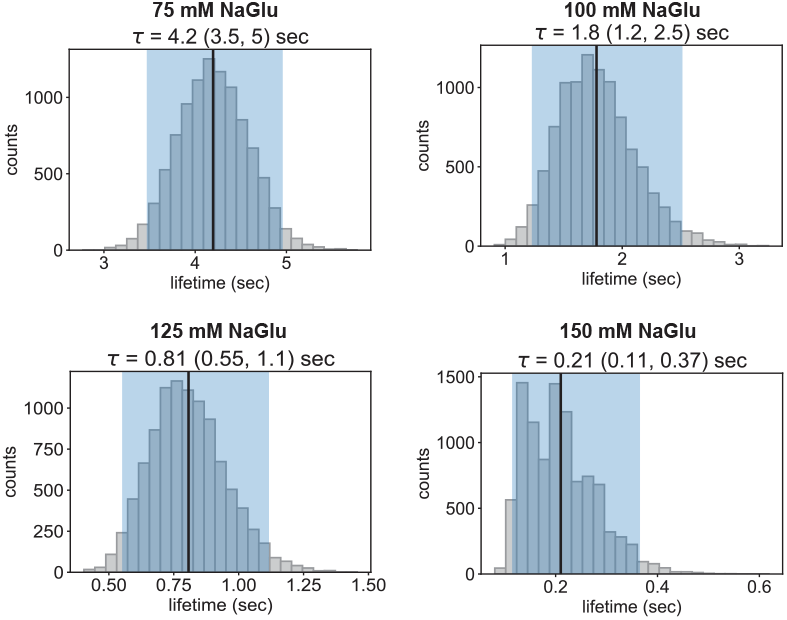
Dwelltime bootstrapping distributions. For each NaGlu concentration, we generated a distribution of dwell times by applying 10000 iterations of a bootstrapping procedure. This approach allowed us to estimate the statistical variability of the measurements and to determine the confidence intervals associated with the mean dwell-time values.

### DNA relaxation assay

In Fig.4 is displayed the relaxation experiment performed using topoII-ΔCTD varying NaCl in independent triplicates. 200 ng of pBR322 (NEB) were incubated with 34.5 ng of Topo2 with in a 10 ul volume containing 50mM Tris–HCl (pH 7.5), 25 mM KCl, 5 mM MgCl2, 1 mM. ATP was added to 1mM and the mixture was incubated at 37 C during 30 min. Reactions were stopped with the addition of 20mM EDTA, 1% (w/v) sodium dodecyl sulphate (SDS) and 100ng/ml proteinase K, and incubated for 1 h at 60 C. Reaction samples were loaded onto 1% (w/v) agarose gels. DNA electrophoresis was carried out at 1.3 V/cm for 15h in TBE buffer (50mM Tris-borate, 1mM EDTA) containing 0.6 ug/ml chloroquine. Gels were stained with Sybr Gold (Thermo Fisher Scientific), destained in water and photographed over an ultraviolet light source. Plots of Lk distributions and quantification of Lk topoisomers were done using ImageJ/Fiji.

### DNA decatenation assay and analysis

In Fig.5 is presented the decatenation experiment performed using topoII-ΔCTD varying NaCl in independent triplicates. 200 ng of Kinetoplast DNA (kDNA, Inspiralis) were incubated with 34.5 ng of Topo2 with in a 10 *µ*l volume containing 50mM Tris–HCl (pH 7.5), 25 mM KCl, 5 mM MgCl2, 1 mM. ATP was added to 1mM and the mixture was incubated at 37 °C during 30 min. Reactions were stopped with the addition of 20mM EDTA, 1% (w/v) sodium dodecyl sulphate (SDS) and 100ng/ml proteinase K, and incubated for 1 h at 60 °C. Reaction samples were loaded onto 1% (w/v) agarose gels. DNA electrophoresis was carried out at 2 V/cm for 15h in TBE buffer (50mM Tris-borate, 1mM EDTA). Gels were stained with Sybr Gold (Thermo Fisher Scientific), destained in water and and photographed over an ultra-violet light source. Plots of the fraction of decatenated minicircles from kDNA were obtained by analysing the gels using ImageJ/Fiji. The analysis was carried out by despeckling the images, and background was subtracted to ensure uniform noise across both axes. Band intensities were manually selected based on local intensity minima surrounding each band. Background noise was estimated from five randomly chosen regions lacking signal and subtracted from each measurement. To account for pipetting variability, band intensities within each lane were normalized to one. Quantification was based on the average of three independent replicates run on the same gel.

### Monovalent Cation Dependence of Relaxation and Decatenation by Full-Length TopoII–Atto647

To ensure that the opposite trends observed for relaxation and decatenation, were not specific to NaCl or to the ΔCTD-topo II, we repeated both assays using NaGlu and full-length topo II labeled with Atto647, in conditions matching those of the optical-tweezers experiments. As shown in Fig. 6a,c, the distribution of super-coils is broader (less efficient) for higher cation concentrations. The opposite effect is observable in Fig. 6b,d, where higher salt concentration leads to more efficient decatenation. Overall, the same qualitative behaviour is recovered, demonstrating that the effect is robust with respect to the salt types and the presence of the CTD.

### Molecular Dynamics Simulations of DNA Relaxation

DNA is modelled as a twistable elastic chain [2, 3]. Briefly, the backbone is made of *N* = 200 beads, each representing 2.5 nm or around 7 bp of DNA (total length *N* = 1500 bp), and decorated by three patches which have no steric interaction with each other. The backbone beads interact via a purely repulsive Lennard-Jones potential as

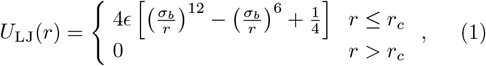

where *r* denotes the separation between the bead centers. The cutoff distance *r*_*c*_ = 2^1*/*6^*σ* is chosen so that only the repulsive part of the Lennard-Jones is used. The energy scale is set by *ϵ* = *κ*_*B*_*T* and the length scale by *σ*_*b*_, both of which are set to unity in our simulations (in this work all quantities are reported in reduced LJ units). Nearest-neighbour beads along the backbone are connected by finitely extensible nonlinear elastic (FENE) springs as

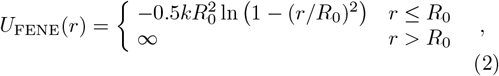

where 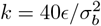 is the spring constant and *R*_0_ = 1.6*σ*_*b*_ is the maximum extension of the elastic FENE bond. To capture DNA’s stiffness (*l*_*p*_ = 150 bp= 50 nm) we introduce a bending energy penalty between consecutive triplets of neighbouring beads along the backbone as

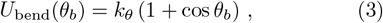

where *θ*_*b*_ is the angle formed between adjacent bonds, i.e. ***t***_*i*_·***t***_*i*+1_*/*|***t***_*i*_||***t***_*i*+1_| with ***t***_*i*_ the tangent at *i*, and *k*_*θ*_ = 20*κ*_*B*_*T* is the bending constant. With this choice *l*_*p*_ = 20*σ*_*b*_ ≃ 50 nm is the persistence length. To capture torsional stiffness, two dihedral CHARMM springs constrain the relative rotation of consecutive beads, *ψ*, at a user-defined value (*ψ*_0_). The torsional angle *ψ* is determined as the angle between planes defined by the triplets bead-bead-patch running along the DNA backbone. The potential

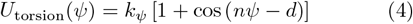

where *k*_*ψ*_ = 50*k*_*B*_*T, n* = 1 and *d* = *ψ*_0_ constrains the rotation. The angle *ψ*_0_ directly determines the thermo-dynamically preferred pitch of the twisted polymer as *p* = 2*π/ψ*_0_ and, in turn, it determines the preferred linking number as *Lk* = *N/p*. In this model, we thus define the supercoiling as *σ ≡ Lk/M* = 1/*p*, which is set by initialising the patchy-polymer as a flat ribbon and by subsequently imposing the angle *ψ*_0_ in order to achieve the wanted *σ* (which may be zero, if *ψ*_0_ = 0 or *p* = ∞). Finally, to maintain consecutive beads parallel to the backbone, we constrain the angle between the triplets bead-bead-patch to *π/*2 so that the frames of reference formed by the triplets are aligned to each other. This potential is written as

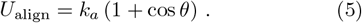

where *θ* is the tilt angle (see Fig of main text) and with *k*_*a*_ = 200*k*_*B*_*T*. The simulations are performed by evolving the equation of motion for the beads coupled to a heat bath which provides noise and friction. The equation of motion for each Cartesian component is thus given by

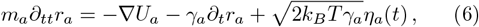

where *m*_*a*_ and *γ*_*a*_ are the mass and the friction coefficient of bead *a*, and *η*_*a*_ is its stochastic noise vector satisfying the fluctuation-dissipation theorem. *U* is the sum of the energy fields described above. The simulations are performed in LAMMPS [4] with *m* = *γ* = *k*_*B*_ = *T* = 1 and using a velocity-Verlet algorithm with integration time step Δ*t* = 0.01 *τ*_*B*_, where *τ*_*B*_ = *γσ*^2^*/k*_*B*_*T* ≃ 0.03 *µ*s (using *γ* = 3*πη*_*water*_*σ* with *η*_*water*_ = 1 cP and *σ* = 2.5 nm) is the Brownian time. See https://git.ecdf.ed.ac.uk/dmichiel/supercoiledplasmids/ for sample codes to reproduce these simulations.

We have chosen the length of the topo II-bound segment to be 5 beads ≃12.5 nm and is modelled by imposing fully phantom interactions with the other beads. The position of this “phantom” segment is then updated as follows: every Brownian time we draw a random number between 0 and 1 and if this is smaller than the dissociation (unbinding) rate *k*_*off*_, the phantom segment is temporarily turned into one that has semi-soft interactions (energy barrier *A* = 10*k*_*B*_*T*) with the other beads in the polymer, and finally after another Brownian time, turned into a fully steric (WCA interaction) segment. The transient semi-soft segment is used to avoid numerical instabilities. If at a particular timestep there is no Topo bound to the polymer, we stochastically introduce one bound Topo at a rate *k*_*on*_. Increasing NaCl in solution is equivalent to increasing *k*_*off*_ (or decreasing the residency time *τ*_*off*_).

We simulate the relaxation for different values of *k*_*off*_ and starting from plasmids at supercoiling value *σ* = 6%. In our model the twist is mostly fixed (due to the large stiffness of the associated dihedrals) and we therefore ex-pect the writhe to change in time. We measure the time-dependence of the writhe by computing the Gauss linking integral [5, 6]

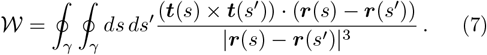

The averages of the relaxation process are performed over at least 50 independent replicas, and for each of these replicas we pre-equilibrate the chains at fixed topology (i.e., without topo II) for at least one relaxation time of the chain, i.e. 10^3^ *τ*_*B*_, where *τ*_*B*_ is the Brownian time.

The molecular dynamics are performed in implicit solvent (Langevin) and are evolved within LAMMPS [4], coupled to an external C++ code that performs the dynamic update of the Topo region. (We provide sample codes in the GitLab repository https://git.ecdf.ed.ac.uk/taplab).

**FIG. 4.**
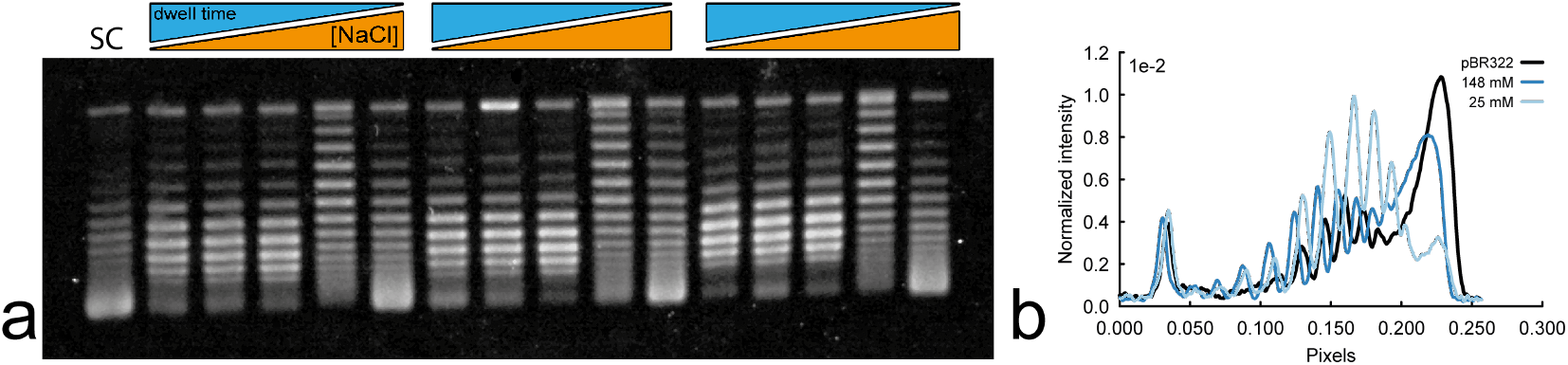
Gel to quantify pBR322 relaxation by topo II. **a**. Gel electrophoresis of 20 ng/ul pBR322 after 30 min incubation with 3.45 ng/ul topoisomerase II. Lane 1 shows supercoiled (SC) pBR322 plasmid; subsequent lanes show three independent replicates at NaCl concentrations of 25, 48, 87, 120, and 148 mM, run on the same gel. **b**. Normalized intensity profiles showing the different supercoil distributions for pBR322, 25mM and 148mM NaCl concentration.

**FIG. 5.**
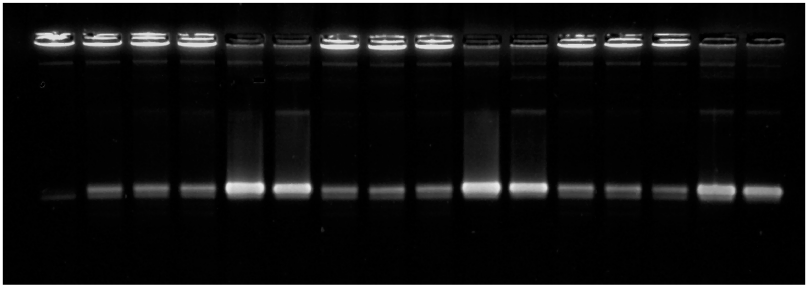
Gel to quantify kDNA decatenation by topo II. Gel electrophoresis of 20 ng/ul kDNA after 30 min incubation with 3.45 ng/ul topoisomerase II. Lane 1 shows catenated kinetoplast DNA; subsequent lanes show three independent replicates at NaCl concentrations of 25, 48, 87, 120, and 148 mM, run on the same gel.

**FIG. 6.**
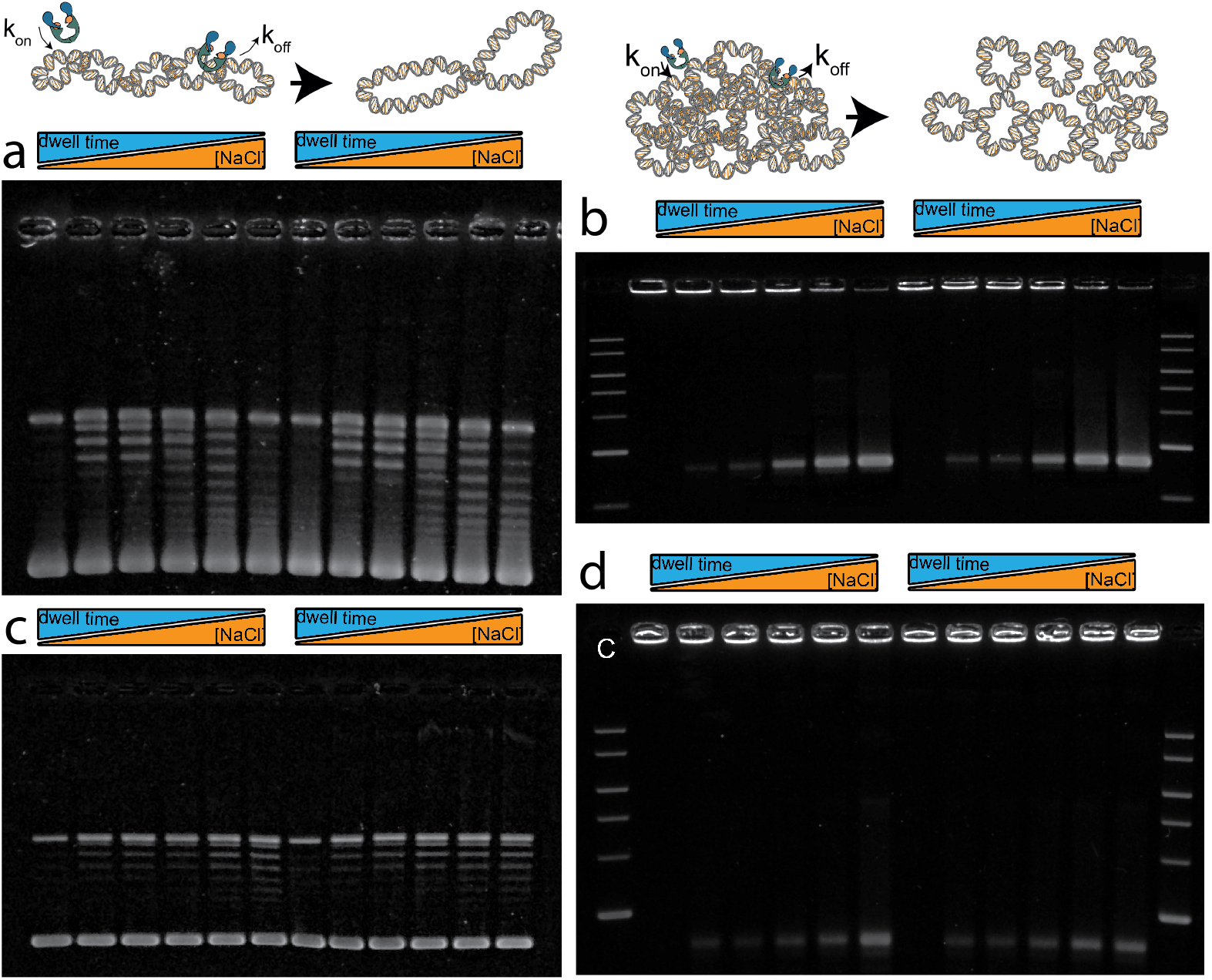
Gels showing relaxation and decatenation by full-length topo II in presence of NaCl and NaGlu. **a**. Gel electrophoresis of 20 ng/ul pBR322 after 30 min incubation with 3.45 ng/ul full-length topo II in presence of NaCl. Lane 1 shows supercoiled (SC) pBR322 plasmid; subsequent lanes show three independent replicates at NaCl concentrations of 25, 48, 87, 120, and 148 mM, run on the same gel. **b**. Gel electrophoresis of 20 ng/ul kDNA after 30 minutes incubation with 3.45 ng/ul of full-length topo II in presence of NaCl. The first lane shows the control sample without protein, and the following five lanes correspond to reactions performed in buffers containing [NaCl] = 25, 48, 87, 120, and 148 mM, respectively. **c**. Gel electrophoresis of 20 ng/ul pBR322 after 30 min incubation with 3.45 ng/ul full-length topoII-atto647 in presence of NaGlu. Lane 1 shows supercoiled (SC) pBR322 plasmid; subsequent lanes show three independent replicates at NaGlu concentrations of 25, 48, 87, 120, and 148 mM, run on the same gel.**d**. Gel electrophoresis of 20 ng/ul kDNA after 30 minutes incubation with 3.45 ng/ul of full-length topoII-atto647 in presence of NaGlu. The first lane shows the control sample without protein, and the following five lanes correspond to reactions performed in buffers containing [NaGlu] = 25, 48, 87, 120, and 148 mM, respectively.

**FIG. 7.**
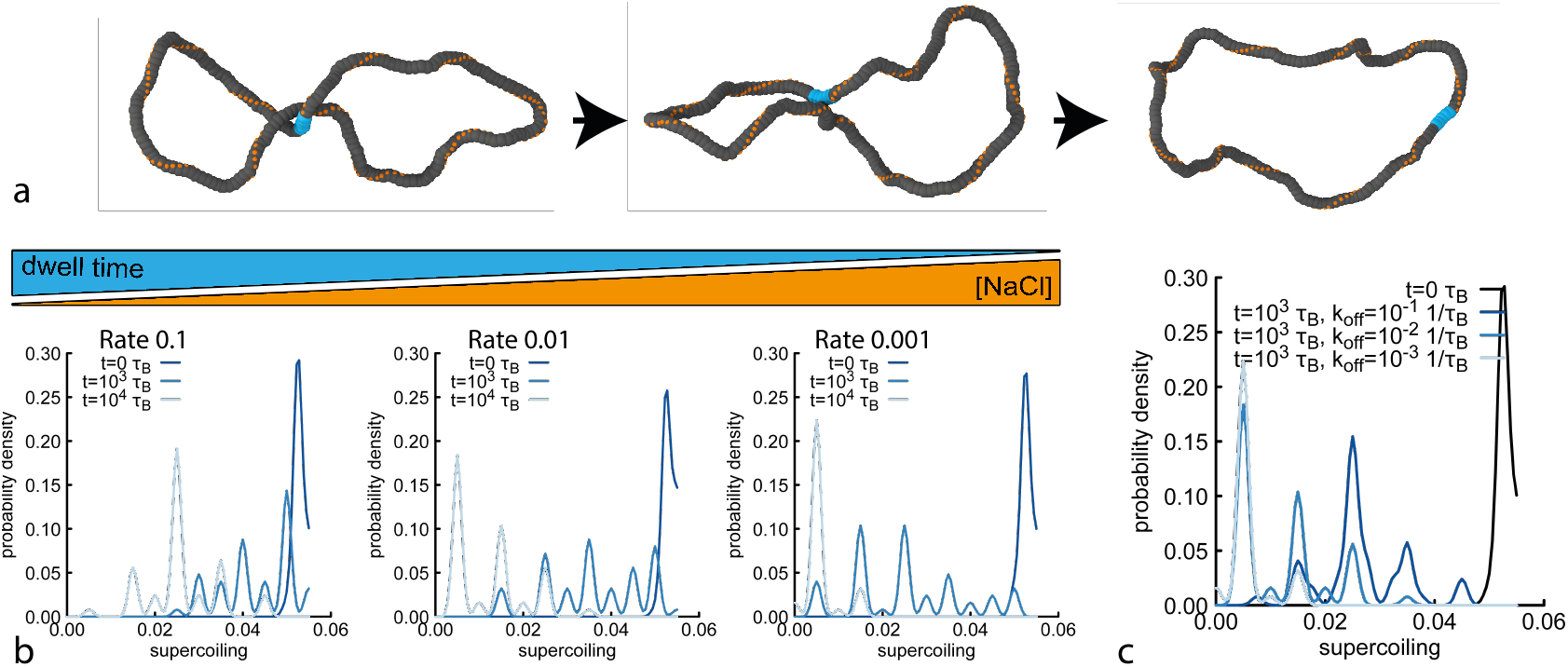
Simulations of DNA plasmids. **a**. Snapshot capturing a simulated event where a crossing induced by supercoiling is simplified by topo II. **b**. Probability density of supercoil distributions at different simulation times, shown for decreasing dwell times from left to right. **c**. Probability density of supercoil distributions three different dwell times compared to the initial distribution.

### Molecular Dynamics of Decatenation

Similarly to what discussed in the previous section, we simulated the molecular dynamics of linked plasmids whose backbone is made of 85 beads. Our initial configurations contains 90 of these plasmids, averaging 3 links per ring. The initial configuration is taken from the MD simulation of a 1 *µ*m^2^ patch of real kDNA from Ref. [7] which forms a connected percolating component, i.e. all rings are linked with at least one other ring. The action of Topo is modelled as a dynamically appearing/disappearing phantom segment, leading to a progressive unlinking of the rings due to strand crossing. In these simulations we apply sub-stoichiometric amounts of topo II (on average 0.86 per ring). Since these simulations are more sensitive to numerical instabilities, the transient semi-soft segment is conserved for 10 Brownian Times instead of 1 as before. To quantify the degree of unlinking we compute the Gaussian linking number between each pair of plasmids in the system, i.e.

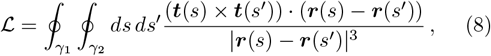

where *γ*_1_ and *γ*_2_ now represent two different plasmids. We thus compute the decatenated fraction as a function of time as the ratio between the number of plasmids with ℒ*k* = 0 at a given time divided by the total number of plasmids at the beginning of the simulation (they are all linked in one percolating component).

**FIG. 8.**
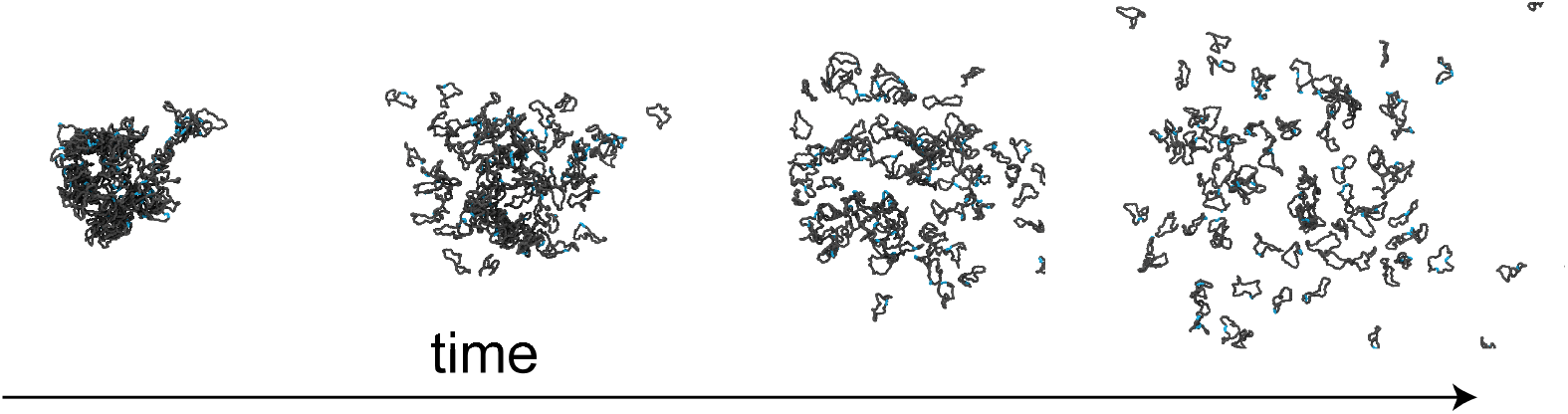
Simulations of kDNA decatenation. Snapshots of catenated DNA structures before topoisomerase II activity and their progressive decatenation over time.

